# Pyrazinamide triggers degradation of its target aspartate decarboxylase

**DOI:** 10.1101/674416

**Authors:** Pooja Gopal, Jickky Sarathy, Michelle Yee, Priya Ragunathan, Joon Shin, Shashi Bhushan, Junhao Zhu, Tatos Akopian, Olga Kandror, Teck Kwang Lim, Martin Gengenbacher, Qingsong Lin, Eric J. Rubin, Gerhard Grüber, Thomas Dick

## Abstract

The introduction of pyrazinamide (PZA) in the tuberculosis drug regimen shortened treatment from 12 to 6 months ^1^. PZA is a prodrug that is activated by a *Mycobacterium tuberculosis* (Mtb) amidase to release its bioactive component pyrazinoic acid (POA) ^2^. Aspartate decarboxylase PanD, a proenzyme activated by autocatalytic cleavage (Supplementary Fig. 1A, ^3^) and required for Coenzyme A (CoA) biosynthesis, emerged as a target of POA ^4-7^. *In vitro* and *in vivo* screening to isolate spontaneous POA-resistant Mtb mutants identified missense mutations in either *panD* or the unfoldase *clpC1*, encoding a component of the caseinolytic protease ClpC1-ClpP ^4,6-9^. Overexpression and binding studies of PanD or ClpC1 pointed to PanD as the direct target of POA whereas *clpC1* mutations appeared to indirectly cause resistance ^4,5,7,9,10^. Indeed, supplementing growth media with CoA precursors downstream of the PanD catalyzed step conferred POA resistance ^4,7,11^. Metabolomic analyses and biophysical studies using recombinant proteins confirmed targeting of PanD by POA ^5^. However, the exact molecular mechanism of PanD inhibition by POA remained unknown. While most drugs act by inhibiting protein function upon target binding, we show here that POA is not a *bona fide* enzyme inhibitor. Rather, POA binding to PanD triggers degradation of the protein by ClpC1-ClpP. Thus, the old tuberculosis drug PZA promotes degradation of its target. While novel for an antibacterial, drug-induced target degradation has recently emerged as a strategy in drug discovery across disease indications. Our findings provide the basis for the rational discovery of next generation PZA.

## Results and Discussion

To characterize POA’s on-target activity, we measured the inhibitory effect of the drug on the enzymatic conversion of aspartate to β-alanine by PanD. Surprisingly, we only observed a weak effect at high concentrations. Although POA binds PanD with a dissociation constant KD of 6 µM ^5^, 200 µM POA reduced β-alanine production by only 15% (Supplementary Fig. 2). Even ‘bathing’ the enzyme in 2 mM POA reduced product formation by 50% only (Supplementary Fig. 2). These weak effects were not significantly attenuated when a POA-resistant PanD mutant that does not bind POA was used (Supplementary Fig. 2). This suggests that POA is not a *bona fide* inhibitor of PanD enzymatic activity.

If POA does not inhibit the catalytic activity of PanD, how does it block the PanD-catalyzed step in CoA synthesis? Interestingly, mutations in PanD and ClpC1 cause the *same* level of PZA/POA resistance in Mtb ^8^. This suggests a mechanistic link between these two proteins and POA. The unfoldase ClpC1 is part of the caseinolytic protease complex ClpC1-ClpP, involved in degradation of substrate proteins ^12,13^. In ClpC1, POA resistance mutations are found primarily in the N-terminal and middle domains, proposed to affect substrate selectivity of the complex ^8,10^. Thus, we speculated that PanD may be recognized and degraded by this machinery. ClpC1-ClpP recognizes substrates via short C-terminal tags ^12-14^. Curiously, mycobacterial PanD proteins contain a 13 amino acid C-terminal extension of unknown function ^15^. Mutations in this tail cause POA resistance and prevent drug binding ^4-7^. Therefore, we hypothesized that (i) the C-terminal tail of Mtb PanD constitutes a degradation tag that is recognized by ClpC1-ClpP and (ii) binding of POA to PanD triggers increased degradation of the target.

To test these hypotheses, we constructed red fluorescent protein (RFP) C-terminal reporter fusions ^14^ and measured the effect of various PanD modifications on protein levels in Mtb (Fig. 1A). As expected, expression of native RFP alone showed a high fluorescence baseline (Fig. 1B). Attachment of PanD’s 13-amino acid C-terminal tail to RFP caused a reduction in fluorescence, indicating that the tail acts as a degradation signal. The same effect was observed when full length PanD was fused to RFP. Consistent with the C-terminal tail acting as a degradation tag, fusion of PanD lacking its native C-terminal tail to RFP restored baseline fluorescence levels. Similar fluorescence was also observed for a fusion carrying a missense mutation in the C-terminal tail ^7^, consistent with amino acid-sequence recognition by the degradation machinery (Fig. 1B). These experiments were repeated independently in *M. bovis* BCG, yielding the same results (Supplementary Fig. 4). To determine whether ClpC1 is involved in degradation of PanD’s C-terminal tail, we measured the effect of introducing each RFP fusion into a POA-resistant *clpC1* mutant strain (Fig. 1B). The level of native RFP was not affected by the *clpC1* mutation compared to wild-type *clpC1*, and baseline fluorescence levels were observed in both genetic backgrounds. As hypothesized, RFP fusions containing the C-terminal tail alone or full length PanD showed increased fluorescence levels in *clpC1* mutant background, indicating impaired degradation activity by the POA-resistant *clpC1* mutant. These results suggested that PanD’s C-terminal degradation tag is recognized by ClpC1, which led us to postulate that PanD degradation is mediated by the caseinolytic protease ClpP. As *clpP* is genetically essential and cannot be deleted, we employed a pharmacological approach. Treatment of Mtb RFP-fusion strains with bortezomib, a mycobacterial ClpP inhibitor ^16^, increased fluorescence of cultures containing RFP-C-terminal tail as well as RFP-full length PanD fusions to the baseline levels of native RFP (Fig. 1C). We further confirmed ClpC1-ClpP mediated recognition and degradation of PanD by *in vitro* reconstitution of the process using recombinant ClpC1 and ClpP proteins ^17^, and cell-free synthesized N-terminally eGFP-tagged PanD as substrate (Supplementary Fig. 5). Taken together, these data demonstrate that PanD is a substrate of the caseinolytic protease complex ClpC1–ClpP and that the mycobacterium-specific tail of PanD is a tag recognized by the protease complex. Our finding is consistent with a previous screen for substrates of ClpP, in which Raju *et al.* identified PanD as one of the 132 Mtb proteins over-represented upon conditional depletion of Mtb ClpP ^13^.

**Figure 1.**
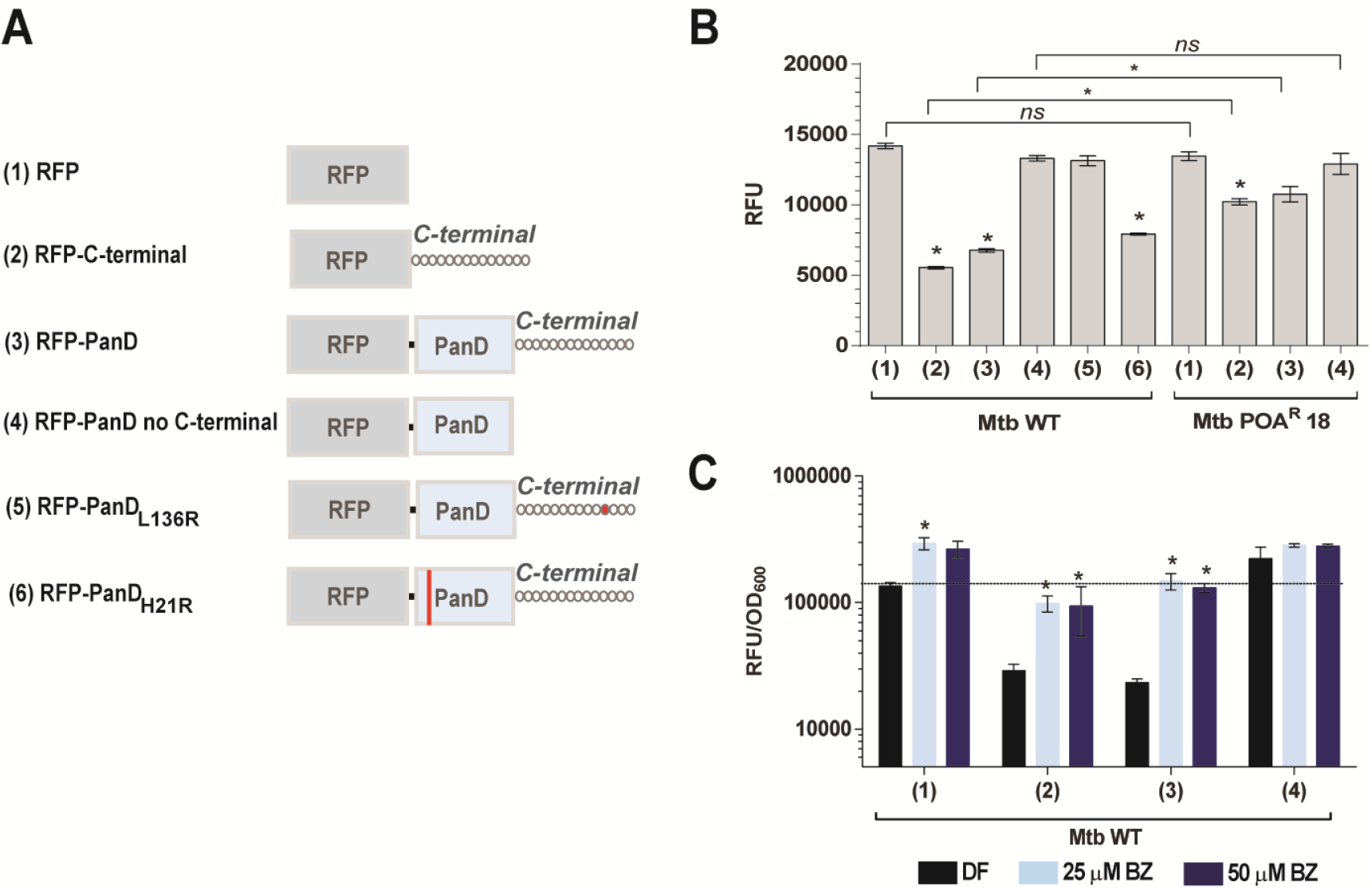
Effect of fusion of various PanD derivatives on the level of RFP reporter protein. (A) Schematic of derivatives of PanD fused translationally to constitutively expressed red fluorescence protein (RFP) as episomal reporter on plasmid pMV262 and transformed into Mtb. “C-terminal” = mycobacterium-specific C-terminal tail of PanD. RFP-PanDL136R and RFP-PanD_H_21R are PZA/POA resistant mutants harboring missense mutation in the N-terminal His21 or the C-terminal tail Leu136 (indicated in red) and have been described previously ^5,7^. (B) Fluorescence levels (expressed as RFU, relative fluorescence units) of mid-log phase cultures (OD_600_= 0.2) were determined as a measure of intra-bacterial fusion protein level in either Mtb wild-type (WT) or PZA/POA-resistant mutant Mtb POA^R^18 [*clpC1*: Lys209Glu] ^8^ harboring RFP-PanD reporter constructs shown in (A). One-way ANOVA multiple comparisons test (Dunn’s posttest, GraphPad Prism) was used to compare fluorescence levels conferred by the various constructs relative to the native RFP in wild-type background (left cluster 1 to 6) and in *clpC1* mutant background (right cluster 1 to 4) respectively; asterisks (*) indicate *p*-value < 0.05; ns: *p*-value > 0.05. The Mann-Whitney test (GraphPad Prism) was used to compare fluorescence levels conferred by each RFP construct in the *clpC1* mutant versus wild-type background (horizontal bars placed across left and right clusters). Asterisks (*) indicate *p*-value < 0.05; ns: *p*-value > 0.05. (C) Effect of bortezomib (BZ) on fluorescence of Mtb cultures carrying different RFP-PanD reporter constructs shown in (A). Cultures were treated with bortezomib (Mtb MIC_50_ = 25 µM) for 3 days. One-way ANOVA multiple comparisons and Dunn’s posttest (GraphPad Prism) was used to compare fluorescence levels in drug-free versus BZ-treated cultures. (*): *p*-value < 0.05. The dotted line indicates mean normalized culture fluorescence of drug-free RFP controls. Experiments were repeated 3 times independently with technical replicates. Means and standard deviations from biological replicate experiments are shown.

Having established that PanD’s C-terminal tail is a degradation tag and PanD levels are regulated post-translationally, we asked whether POA treatment enhances PanD degradation. POA treatment of cultures harboring the native RFP did not affect fluorescence (Fig. 2A), neither did it affect fluorescence of cultures expressing the RFP-C-terminal fusion, consistent with the requirement of a dual epitope (including His21, Fig. 1A) for POA binding to PanD ^5^. In contrast, the full length PanD fusion showed a dose-dependent reduction of fluorescence by POA. This reduction was dependent on PanD’s C-terminal tail and was not observed in cultures harboring fusion with POA-resistant PanD proteins containing missense mutations in either the N-terminal His21 or the C-terminal tail, both abolishing binding of POA (Fig. 2A, ^5^). To confirm that POA indeed causes reduction of endogenous PanD levels, we performed Western blotting with an anti-PanD antibody, and showed rapid reduction of native PanD in response to POA treatment in wild-type *M. bovis* BCG but not in a POA-resistant C-terminal PanD missense mutant (Supplementary Fig. 6, 7). Further, RFP reporter-based work showed that increased degradation of PanD in Mtb was also observed upon treatment with PZA, the prodrug of POA (Fig. 2A), but not with the control drug isoniazid (Fig. 2B), that targets mycolic acid synthesis. This showed that increased PanD degradation is POA/PZA-specific and not a general drug treatment effect. PanD mRNA levels were not affected by POA treatment, ruling out transcriptional effects (Supplementary Fig. 8). To determine whether ClpC1 is involved in POA’s mechanism of action, we tested the impact of *clpC1* mutation on POA-induced PanD degradation by measuring the effect of POA on the various RFP-PanD fusions in the *clpC1* mutant background. POA-resistant *clpC1* mutations prevented POA/PZA-induced reduction of PanD levels (Fig. 2C). To determine whether the ClpP protease is required for POA-induced PanD degradation, we co-treated cultures with POA and the ClpP inhibitor bortezomib. Inhibition of ClpP prevented POA-induced reduction of PanD levels (Fig. 2D). It is to note that bortezomib also increased fluorescence of the RFP control strain, although to a lesser extent than the PanD-RFP reporter strains. The reasons for this effect remain to be investigated and may include yet unidentified off-target effects. To determine whether the other major protein degradation machinery of Mtb, the proteasome ^18^, contributes to PanD degradation, we measured the effect of *prcAB* gene deletion on POA susceptibility. POA MIC was not affected by the deletion, arguing against proteasome involvement in PanD degradation (Supplementary Fig. 9). Together, these results suggest that the mechanism of action of PZA/POA involves suicidal derailing of a bacterial post-translational regulatory mechanism in which POA promotes the breakdown of its essential target by Mtb.

**Figure 2.**
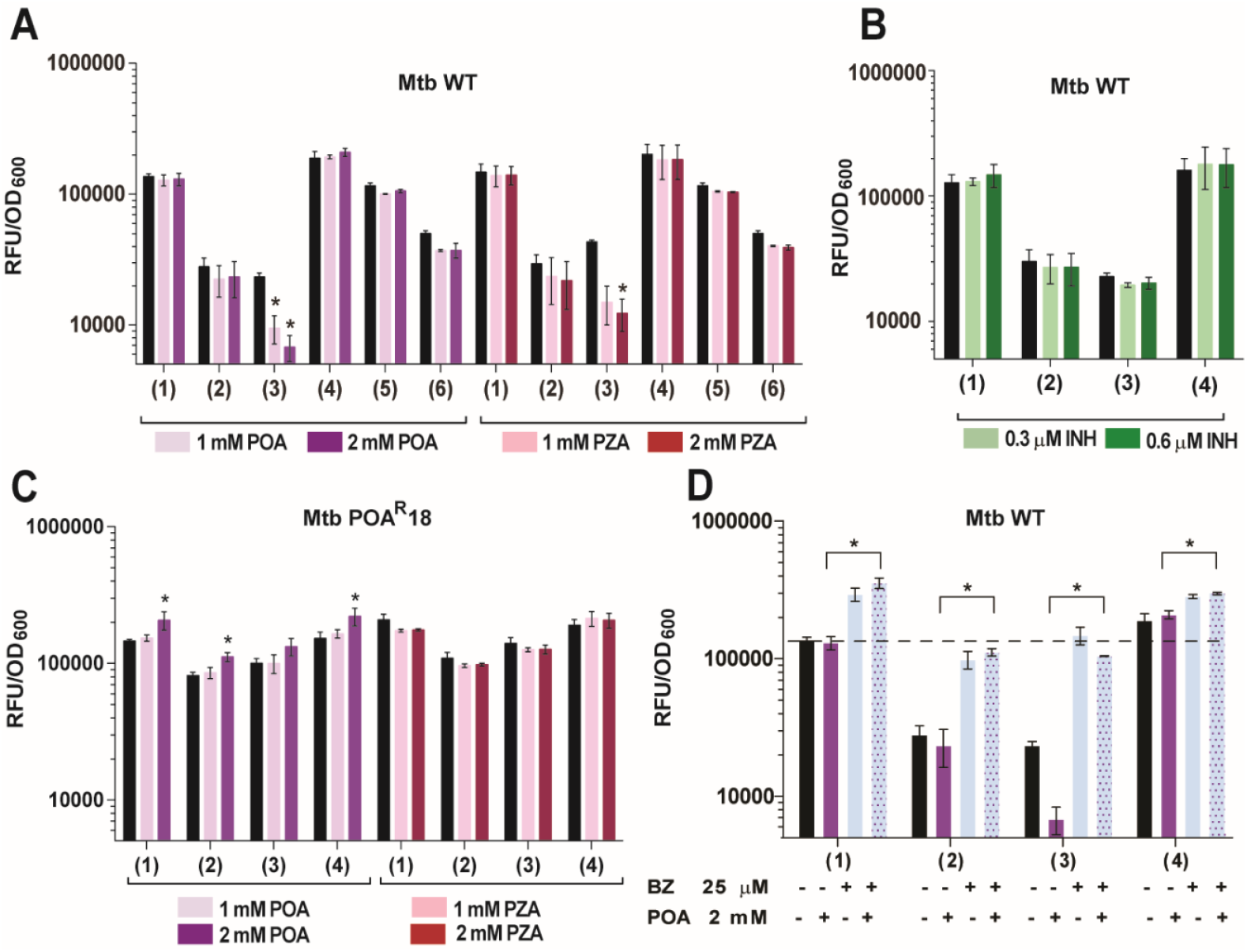
Effect of POA, PZA, INH treatment and POA/bortezomib co-treatment on levels of various RFP-PanD fusions. Mtb strains harboring various RFP-PanD fusions shown in Fig. 1A were treated with increasing doses of POA, bortezomib (BZ) and isoniazid as control drug (INH) for 3 days. Fluorescence (RFU) was measured and normalized to bacterial growth (OD_600_). (A) RFP-PanD fusions in Mtb wild type (WT) treated with POA (1 mM, or Mtb MIC_50_) and PZA. (B) RFP-PanD fusions in Mtb wild type treated with isoniazid (INH): 0.3 µM, or Mtb MIC_50_. (C) RFP-PanD fusions in POA-resistant *clpC1* mutant Mtb POA^R^18 [*clpC1*: Lys209Glu] treated with POA and PZA. (D) RFP-PanD fusions in Mtb wild-type treated with BZ (25 µM, or Mtb MIC_50_) or POA (2 mM) alone or in combination as indicated. The dashed line indicates mean normalized culture fluorescence of drug-free RFP controls. (A, B, C) One-way ANOVA multiple comparisons test (Dunn’s posttest, GraphPad Prism) was used to compare fluorescence levels in drug-treated versus drug-free controls. (*) indicate *p*-value < 0.05. (D) The Mann-Whitney *U* test (GraphPad Prism) was used to compare the effect of POA treatment on fluorescence levels conferred by RFP constructs 1 to 4 (wild-type background, Fig. 1A) in the presence and absence of BZ. (*) indicate *p*-value < 0.05. For each strain and treatment, drug-free controls were included (black bars). Experiments were carried out three times independently. Means and standard deviations from biological replicate experiments are shown.

In the absence of POA, PanD’s C-terminal tail may be poorly accessible, resulting in low degradation efficiency. Binding of POA may induce structural changes of PanD, rendering the protein’s degradation tag more accessible to the protease complex, thus allowing increased degradation. Indeed, dynamic light scattering experiments (Supplementary Fig. 10), electron microscopy (Supplementary Fig. 11) and circular dichroism (Supplementary Fig. 12) analyses revealed that POA has dramatic effects on the quaternary and secondary structures of wild-type PanD but not on the POA resistant (non-POA binding) mutant protein. The precise structural mechanism of POA-mediated increased degradation of PanD by ClpC1-ClpP remains to be determined.

Drug-induced degradation of a target protein is a novel antibacterial mechanism. However, this strategy does have precedents: selective estrogen receptor degraders (SERDs) such as Fulvestrant, used clinically for the treatment of certain breast cancers, are known to decrease intracellular estrogen receptor α levels ^19^. In addition to Fulvestrant, small-molecule phthalimides (e.g. Thalidomide and Lenalidomide) ^20,21^ and the plant hormone Auxin ^22^ were shown to induce the degradation of specific substrates. Interestingly, targeted protein degradation as a novel drug discovery approach has gained momentum in recent years ^23^. PROTACs, heterobifunctional molecules which contain discrete binding moieties for the protein of interest and for E3 ligase, make use of the human ubiquitin – proteasome system to specifically degrade tagged proteins ^24^. A first PROTAC drug is heading for clinical trials ^25^. In conclusion, we report that an old tuberculosis drug acts by an event-based rather than occupation-based mechanism, accelerating degradation of its cellular target by Mtb’s own protease machinery.

## Methods

### Bacterial strains, culture medium and chemicals

*M. tuberculosis* H37Rv (ATCC 27294) and *M. bovis* BCG (ATCC 35734) strains were maintained in complete Middlebrook 7H9 medium (BD Difco) supplemented with 0.05% (vol / vol) Tween 80, 0.5% (vol / vol) glycerol, 0.5% albumin, 0.2% glucose, 0.085% sodium chloride and 0.0003% catalase at 310 K with agitation at 80 rpm. POA-resistant strain *M. tuberculosis* H37Rv POA^R^18 (*clpC1*: A625G/Lys209Glu) was isolated and described in ^8^. The *M. tuberculosis* H37Rv Δ*prcAB* strain (Hygromycin-resistant) and its complemented mutant (Hygromycin-resistant and Kanamycin-resistant) were generated and described in ^17^. Hygromycin B (Roche) and Kanamycin (Sigma-Aldrich) were used for selection when required at 50 µg/mL and 25 µg/mL respectively. Pyrazinamide, Pyrazinoic acid and Isoniazid were purchased from Sigma-Aldrich. Bortezomib was purchased from Chembridge. Antibiotics were dissolved in 90% DMSO and sterilized using 0.2 μm PTFE membrane filters (Acrodisc PALL).

### Susceptibility testing

MICs were performed by the broth dilution method as described in ^8^. The strains were grown to mid-log phase, spun down, resuspended in fresh 7H9 broth and adjusted to an OD600 = 0.1. 100 µL of cell suspension was added into wells containing 100 µL two-fold serially diluted compound in transparent flat-bottomed 96-well plates (Corning Costar), sealed with Breathe-Easy membranes (Sigma Aldrich). The plates were incubated for 7 days at 310 K with shaking at 80 rpm. After incubation, the cultures were manually resuspended and OD_600_ was measured using a spectrophotometer (Tecan Infinite M200 Pro). Experiments were repeated independently with technical replicates. MIC_50_ values represent the concentration of drug which inhibits bacterial growth by 50% as compared to the respective drug-free control.

### Protein preparation

Wild-type Mtb Aspartate decarboxylase PanD (PanD_WT_) and the POA-resistant mutant PanD (PanD127TRASC131) proteins were overexpressed with N-terminal 6x His tags in *E. coli* and purified by Ni^2+^-NTA affinity, followed by gel filtration chromatography with buffer containing 50 mM Tris-HCl (pH 7.5), 200 mM NaCl as described previously ^*5*^. Protein concentrations were determined using a BioSpec-nano spectrophotometer (Shimadzu, USA). Purity and homogeneity of samples were verified by SDS-PAGE (Supplementary Fig. 1).

### SDS-PAGE analyses of recombinant PanD

Protein samples were analyzed by 12% SDS-PAGE (NuPAGE Bis-Tris precast gel) run at a constant voltage of 200 V for 45 minutes in NuPAGE MES-SDS running buffer. The recombinant protein samples were denatured with a sample buffer containing LDS (NuPAGE), 0.1 M Dithiothreitol (DTT) and heating at 368 K for 10 minutes prior to loading. The gels were stained with Coomassie Brilliant Blue R-250 (BioRad) as described by the suppliers.

### ^**1**^**H-1D NMR studies of L-Asp to β-Ala conversion**

NMR samples containing 2 mM L-Asp, 10 µM of PanD_WT_ or PanD_127TRASC131_ were prepared in deuterated water (D_2_O) buffer. NMR experiments were carried out using a Bruker Avance 400 MHz NMR spectrometer, equipped on a 5 mm BBI probe head at 298 K as described by Sharma *et* al. ^26^. Basic ^1^H-1D NMR spectra were collected with pre-saturation of solvent water (relaxation delay 2 s; solvent pre-saturation applied during the relaxation delay). Time-dependent ^1^H-1D NMR experiments were performed to examine the kinetics of enzymatic conversion of L-Asp to β-Ala at six different incubation time points (0 min, 10 min, 20 min, 30 min, 40 min, 60 min and 90 min) using PanD_WT_ (Supplementary Fig. 2A and 2B). In order to study the inhibitory effect of POA, the compound was added (0.2 and 2 mM) to reaction mixtures, containing 2 mM of L-Asp. 10 µM of enzyme (PanD_WT_ or PanD127TRASC131) was added to initiate the reaction (spectra are shown in Supplementary Fig. 3).

### Fluorescence reporter strains and reporter assay

Genomic DNA was isolated from *M. tuberculosis* H37Rv (ATCC 27294), *M. bovis* BCG (ATCC 35734), POA-resistant strains *M. bovis* BCG POA1.3 (*panD*: A62G/His21Arg), *M. bovis* BCG POA1.1 (*panD*: T407G/Leu136Arg), which were isolated and described in ^7^. The sequences of *panD* in *M. bovis* BCG (ATCC 35734) and *M. tuberculosis* H37Rv (ATCC 27294) are 100% identical as determined by sequencing. Primers and templates used for plasmid construction of each RFP protein fusion are summarized in Supplementary Table 1. PCR amplification was performed with Phusion DNA polymerase (Thermo Scientific) as per manufacturer’s instructions.

The obtained RFP fusion constructs were electroporated into *M. tuberculosis* H37Rv wild-type, *M. tuberculosis* H37Rv POA^R^18 or wild-type *M. bovis* BCG as specified and selected on complete Middlebrook 7H11/7H10 agar containing Kanamycin (25 μg/mL) at 310 K. Kanamycin-resistant single colonies were picked and colony-purified. Strains were grown to log phase in respective growth media and stored in 7H9 broth containing 10% glycerol in 1 mL aliquots at 193 K.

For the reporter assay, the different fluorescent reporter strains were grown to mid-log phase (OD_600_ = 0.4 - 0.6) in their respective selective 7H9 media. All cultures were then adjusted to OD_600_ = 0.4 in fresh 7H9 broth after centrifugation at 3200 rpm for 10 min and 100 µL of this suspension was inoculated into wells of a flat-bottomed 96-well plate (Costar Corning), each containing 100 µL of media (with or without drugs). Reporter assays were carried out with dual readout, absorbance (OD600), and relative fluorescence units (RFU; *λ*ex*/λ*em, 587/630 for RFP) by using an Infinite M200 Pro plate reader (Tecan). Baseline measurements at day 0 were carried out with RFU, after which plates were sealed with Breathe-Easy sealing membranes (Sigma-Aldrich) and incubated at 310 K with shaking at 80 rpm for measurements at subsequent time points. Experiments were carried out with technical replicates in independent biological replicates.

### Molecular cloning and cell-free synthesis of fluorescently-tagged proteins

Constructs encoding eGFP alone, or eGFP fused to the N-terminus of proteins of interest were cloned into pET26b or pET21a to allow T7 polymerase initiated *in vitro* expression. Briefly, expression vectors were linearized by restriction digestion (NdeI and HindIII) and column purified. To make pET26b_eGFP, mycobacterial codon-optimized eGFP was amplified with JH-306 and JH-311, column purified, and cloned into linearized pET26b by Gibson assembly ^27^. To generate eGFP fusion constructs, an 18 bp linker (5’-ggatctagcggatccagt-3’, encoding a flexible GSSGSS linker when being translated) was used as both the DNA homology for Gibson assembly and a flexible linker peptide. PCR amplified eGFP-linker (JH-306/JH-307) and linker-protein-of-interest (Supplementary Table 1) were column purified and cloned into designated vectors using Gibson assembly.

*In vitro* protein expression was carried out following manufacturer’s instructions (NEB, E6800s). Briefly, 300 ng of each column purified plasmid (Supplementary Table 1) was mixed with PURExpress solution A and B (10 and 7.5 µL, respectively), 0.5 µL of RNaseOUT^TM^ RNase inhibitors (Thermo) and nuclease-free water (Promega) to make final volume 25 µL. Protein synthesis was allowed by incubating the reaction mix at 310 K for 5 hours, then stopped by adding excessive amount of chloramphenicol (final concentration as 500 µg/mL). Successful protein production was validated by SDS-PAGE electrophoresis and Coomassie staining. The remaining products were kept at 277 K for no more than a week before further steps.

### *In vitro* reconstitution of PanD degradation

PanD degradation assays with purified recombinant ClpC1 and ClpP (ClpP1 and ClpP2) were carried out as described previously ^17^. Full-length (eGFP_full_PanD) and N-terminally truncated (eGFP_PanD_ΔN-term1-24_, eGFP_PanD_ΔN-term1-24, ΔC-term127-139_) PanD proteins were produced with N-terminally fused eGFP in a cell-free transcription / translation system as described above. Aliquots from protein expression mixes were diluted 5-fold into the proteolysis assay mixture containing 300 nM ClpP1P2 tetra-decamer and 300 nM ClpC1 hexamer in reaction buffer (20 mM K-phosphate buffer pH 7.6 containing 100 mM KCl, 5% glycerol, 8 mM Mg-ATP, and 2.5 mM of dipeptide activator Bz-Leu-Leu). Degradation was followed by the loss of eGFP fluorescence ^17^.

### Quantitative PCR

RNA from *M. tuberculosis H37Rv* wild-type was isolated from an equivalent of 20 mL of culture at an OD_600_ = 0.4. Cultures were spun down at 3200 rpm for 10 min, resuspended in 1 mL TRIzol (Invitrogen), and subjected to bead beating by using a FastPrep-24 5G instrument (MP Biomedicals; twice for 45 s each, 5 min on ice between pulses). RNA was purified using the PureLink RNA mini kit with the Turbo DNA-free kit (Invitrogen) following the manufacturer’s instructions with on-column DNAse treatment (Invitrogen). cDNA was synthesized from 500 ng of total RNA with the SuperScript III first-strand synthesis system (Invitrogen) by using random primers (Promega). Quantitative PCR was performed using the FastStart Essential DNA Green Master (Roche) on a LightCycler 96 real-time PCR system (Roche) (for primers, see Supplementary Table 2). Relative expression of transcripts was determined by the ΔΔC_Q_ method as compared to 16S rRNA which was uniformly expressed in our Mtb strains. cDNA was synthesized from at least two independent RNA samples and qRT-PCR was performed at least twice, in triplicate wells, for each cDNA sample.

### Preparation of mycobacterial cell lysates

*M. bovis* BCG wild-type or mutant cultures were grown to mid-log phase in independent biological replicates and 100 mL of culture (adjusted to OD_600_ = 0.4) was harvested by centrifugation at 3400 x g for 20 min at 277 K, washed with ice-cold phosphate buffered saline and pelleted again. For the Western blotting experiments, mid-log cultures were adjusted to OD_600_ = 0.2 in fresh media and subjected to POA treatment (1 mM or 4 mM), the equivalent of 100 mL OD_600_ = 0.4 was harvested at specified time points. The cell pellets were resuspended in 600 µL lysis buffer (50 mM Tris/Hcl pH 7.5, 5% (vol/vol) glycerol, 1.5 mM MgCl2, 150 mM NaCl, 1mM DTT, 1% n-dodecyl β-D-maltoside (w/vol), 1x complete EDTA-free protease inhibitor cocktail (Roche)), transferred to a lysis matrix B tube (MP Biomedicals) and homogenized using a FastPrep-24 5G instrument (MP Biomedicals). The cell debris was pelleted by centrifugation at 13000 rpm for 10 min at 277 K and the supernatant (about 400 µL) was collected and stored at 193 K until further use. The protein concentration was determined by a BCA protein assay kit (Pierce).

### In-gel digestion

The protein lysates were concentrated by SpeedVac to 30 µL each. Samples were then polymerized with 4% SDS, 10% acrylamide (29:1 C), 0.25% ammonium persulfate and 0.25% TEMED for 30 min. The gels were fixed with 50% methanol, 12% acetic acid for 30 min. The bands of interest were individually cut out from the gel (for protein identification from SDS-PAGE) or the entire polymerized cell lysates (prepared as described above) were cut into ∼1 mm^3^ pieces and washed 3 times with 0.5 mL 50 mM Triethylammonium bicarbonate buffer (TEAB), 50% (vol/vol) acetonitrile and then treated with 500 µL acetonitrile. The gel pieces were reduced with 5 mM tris(2-carboxyethyl)phosphine (TCEP) in 100 mM TEAB and incubated at 329 K for 1 hour. TCEP solution was removed and samples were dehydrated with acetonitrile. Solvent was removed, and samples were alkylated with 10 mM methyl methanethiosulfonate (MMTS) in 100 mM TEAB, followed by incubation at room temperature for 60 min. The gel pieces were washed with 50 mM TEAB and dehydrated with acetonitrile two times. The gel pieces were dried in a Speedvac. 12.5 ng/µL of Trypsin Gold (Promega) in 500 mM TEAB was added and kept at 277 K for 30 min. The mixture was incubated at 310 K for 16 hours. 200 µL 50 mM TEAB was added to the digest, spun down at 6000 rpm for 10 min and the supernatant was collected. 200 µL 5% formic acid (vol/vol), 50% acetonitrile was added to the gel pieces, spun down at 6000 rpm for 10 min and the supernatant was collected. The gel pieces were finally treated with 200 µL acetonitrile, followed by centrifugation at 6000 rpm for 10 min. The three supernatants were combined and dried in a SpeedVac.

### LC–MS/MS analysis

The peptide separation was carried out on an Eksigent nanolC Ultra and ChiPLC-nanoflex (Eksigent, Dublin,CA, USA) in Trap Elute configuration. The samples were desalted with Sep-Pak tC 18 μ Elution Plate (Waters, Milford, MA, USA) and reconstituted with 20 μL of diluent (98% Water, 2% Acetonitrile, 0.05% Formic acid). 5 μL of the sample was loaded on a 200 μm × 0.5 mm trap column and eluted on an analytical 75 μm × 150 mm column. Both trap and analytical columns were made of ChromXP C18-CL, 3 μm (Eksigent, Germany). Peptides were separated by a gradient formed by 2% acetonitrile, 0.1% formic acid (mobile phase A) and 98% acetonitrile, 0.1% formic acid (mobile phase B): 5 to 7% of mobile phase B in 0.1 min, 7 to 30% of mobile phase B in 10 min, 30 to 60% of mobile phase B in 4 min, 60 to 90% of mobile phase B in 1 min, 90 to 90% of mobile phase B in 5 min, 90 to 5% of mobile phase B in 1min and 5 to 5% of mobile phase B in 10 min, at a flow rate of 300 nL/min.

The MS analysis was performed on a TripleTOF 5600 system (AB SCIEX, Foster City, CA, USA) in Information Dependent Mode. MS spectra were acquired across the mass range of 400–1250 m/z in high resolution mode (>30000) using 250 ms accumulation time per spectrum. A maximum of 20 precursors per cycle were chosen for fragmentation from each MS spectrum with 100 ms minimum accumulation time for each precursor and dynamic exclusion for 8 s. Tandem mass spectra were recorded in high sensitivity mode (resolution > 15000) with rolling collision energy. Survey-IDA Experiment, with charge state 2 to 4, which exceeds 125 cps was selected.

Peptide identification was carried out on the ProteinPilot 5.0 software Revision 4769 (AB SCIEX) using the Paragon database search algorithm (5.0.0.4767) for peptide identification and the integrated false discovery rate (FDR) analysis function. The data were searched against a database consisting of *M. bovis* BCG proteome UP000001472 (total 7782 entries). The search parameters are as follows: Sample Type — Identification; Cys Alkylation — MMTS; Digestion — trypsin; Special Factors — None; Species – None. The processing was specified as follows: ID Focus— Biological Modifications; Search Effort — Thorough; Detected Protein Threshold — 0.05 (10.0%) and competitor Error Margin (ProtScore) – 2.00.

### Western blot analyses of native PanD protein

Total protein extracts were prepared from *M. bovis* BCG cultures as described above and protein contents were quantified using the BCA protein assay kit (Pierce). 10 µg of total protein lysates were subjected to 12% SDS PAGE (NuPAGE Bis-Tris precast gel and NuPAGE MES-SDS running buffer). Dual color markers (Biorad Precision Plus) and recombinant PanD_WT_ protein were included as molecular weight markers. Proteins were transferred onto 0.2 µm nitrocellulose membranes (Biorad) in a mini Trans-Blot cell (Biorad). Blots were blocked with 5% skimmed milk containing Phosphate-buffered saline with 0.1% Tween-20 (PBST) for 3 hours at 298 K and then probed with PanD-specific rabbit sera 1:500 in 5% skimmed milk-PBST overnight at 277 K. PanD-specific rabbit sera were obtained from i-DNA Technologies (Singapore) by immunizing rabbits with a synthetic peptide from the PanD aspartate decarboxylase domain (C-IAYATMDDARARTY-amide). Secondary HRP Goat-anti-rabbit antibodies (Invitrogen) were used at a 1:5000 dilution in 5% skimmed milk-PBST for 1 hour at 298 K. Detection was performed with Clarity Max Western ECL kit (Biorad) and visualized with a Chemidoc imaging system (Biorad). To ensure equivalent loading of total protein lysates, blots were stripped and re-probed with Anti-RpoB monoclonal antibody (Abcam) at a 1:10000 dilution in 5% skimmed milk-PBST and 1:20000 secondary HRP Rabbit-anti-mouse antibody (Invitrogen) in 5% skimmed milk-PBST_16_.

### Dynamic light scattering studies

Dynamic light scattering experiments of PanD_WT_ and the mutant PanD_127TRASC131_ in the absence or presence of 200 µM POA were carried out using the Malvern Zetasizer Nano ZS spectrophotometer. The protein was incubated for 40 min with drug before measurement. The samples were measured in a low-volume quartz batch cuvette (ZEN2112, Malvern Instruments) using 12 µL of 1 mg/mL of PanD in 50 mM Tris (pH 7.5) buffer containing 200 mM NaCl. After 60 s equilibration time at 298 K, the backscattering at 173° was detected for all proteins. Scattering intensities were analyzed using the instrument software to calculate the hydrodynamic diameter (D_H_), size, and volume distribution.

### Electron microscopy

PanD_WT_ and the mutant PanD_127TRASC131_ was incubated with 200 μM POA for 40 min as described under Dynamic Light Scattering before applying a volume of 4 μL of the drug-bound protein (50 μg/mL) to a glow discharged carbon-coated copper TEM grid and stained with 2% (v/v) uranyl-acetate. Electron micrographs were recorded on a 120 kV Tecnai spirit T12 transmission electron microscope (FEI) equipped with a 4 K CCD camera (FEI) at a calibrated magnification of 66,350 x under low-dose conditions.

### Circular dichroism spectroscopy

Circular dichroism (CD) spectra of PanD_WT_ and the mutant PanD_127TRASC131_ with or without 200 µM POA were recorded with a CHIRASCAN spectrometer (Applied Photo-physics) using a 60 µL quartz cell (Hellma, Germany) with 0.1 mm path length. The respective protein was incubated for 40 min with drug before measurement. Light of 190-260 nm was used to record the far UV-spectra at 298 K with 1 nm resolution. Two independent measurements were carried out and each measurement was performed three times for each sample. The CD-spectra were acquired in a buffer of 50 mM Tris, pH 7.5 and 200 mM NaCl with protein concentration of 2.0 mg/mL. Secondary structural content was determined from CD spectrum using K2D3 software ^28^.

## Acknowledgments

We thank Sabai Phyu, National University of Singapore BSL3 Core Facility, for support and Uday Ganapathy and Veronique Dartois, Center for Discovery and Innovation - Hackensack Meridian Health, for discussion and comments on the manuscript. We are grateful to Sabine Ehrt, Weill Medical College of Cornell University, for providing the *prcAB* knockout strain. We thank N. Kamariah and S. Rishikesan, School of Biological Sciences, Nanyang Technological University, Singapore, for their support in imaging the PanD proteins, Bill Jacobs, Albert Einstein College of Medicine, for sharing his pMV262 plasmid and Yoshiyuki Yamada, National University of Singapore, for the modified pMV262-mRFP construct. This research was supported by the Singapore Ministry of Health’s National Medical Research Council under NMRC/TCR/011-NUHS/2014 and NMRC/CG/013/2013 (T.D.) and is part of the SPRINT-TB program led by Nick Paton. J.S. received a Yong Loo Lin School of Medicine graduate scholarship. T.D. holds a Toh Chin Chye Visiting Professorship at the National University of Singapore. Research reported in this publication is also supported by the National Institute of Allergy and Infectious Diseases of the National Institutes of Health under Award Numbers 2R01AI106398-05 (T.D.), P01AI095208 (E.J.R), by the Bill and Melinda Gates Foundation OPP1181211 (E.J.R.), and by the National Research Foundation (NRF) Singapore, NRF Competitive Research Programme (CRP), Grant Award Number NRF–CRP18–2017–01; G.G.). The content is solely the responsibility of the authors and does not necessarily represent the official views of the National Institutes of Health.

## Author contributions

Conceptualization, P.G., J.S., M.Y., M.G., E.J.R., G.G., T.D.; Methodology, P.G., J.S., M.Y., M.G., L.Q., E.J.R., G.G. and T.D.; Investigation, P.G., J.S., M.Y., P.R., J.S., S.B., J.Z., T.A., O.K., T.K.L. and M.G.; Writing – Original Draft, P.G., G.G. and T.D.; Writing – Review & Editing, all authors; Funding Acquisition, E.J.R., G.G., T.D.; Supervision, P.G., L.Q., E.J.R., G.G., T.D.

## Author Information

Authors declare no competing interests. Correspondence and requests for materials should be addressed to Thomas Dick.

