## Supplemental Figures and Tables for "Pyrazinamide triggers degradation of its target aspartate decarboxylase"

**Supplementary Data:**

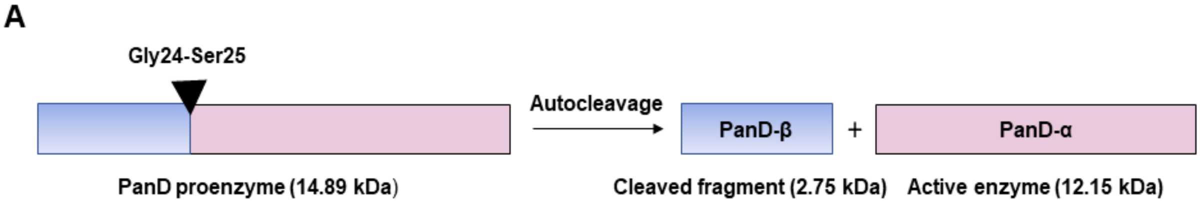


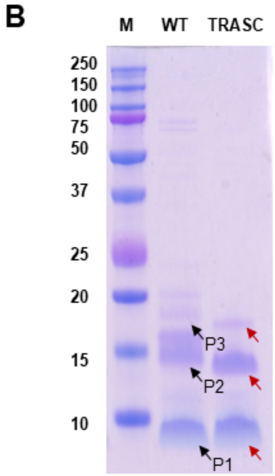


| Calculated molecular weight (kDa) | PanD (WT) | PanD (TRASC) |
|---|---|---|
| Full length proenzyme, including His tag (P3) | 15.9 | 15.15 |
| Activated form (P2) | 12.15 | 11.4 |
| Cleaved N-terminal fragment, including His tag (P1) | 3.75 | 3.75 |

**Supplementary Fig. 1. Verification of purified recombinant PanD proteins.** (A) Schematic depicting the auto-cleavage of PanD at the Gly24-Ser25 position to yield the C-terminal active enzyme (PanD-α) and an N-terminal cleaved fragment (PanD-β) [3]. (B) 12% SDS-PAGE of purified, N-terminal His-tagged recombinant proteins $PanD_{WT}$ (WT) and C-terminal truncated, non-POA binding $PanD_{127TRASC131}$ (TRASC). To determine the identity of the protein bands, the bands corresponding to P1, P2 and P3 were cut out, in-gel digested with trypsin and subjected to LC-MS analysis. PanD is synthesized as a proenzyme and rapidly auto-catalytically cleaved at its N-terminus, resulting in a small N-terminal fragment (P1) and the active C-terminal enzyme (P2). Traces of uncleaved proenzyme are indicated (P3). Black and red arrows indicate the corresponding protein bands of wild-type and truncated PanD, respectively. Note: The cleaved form of the wild-type protein P2 migrates as a doublet with similar staining intensity. The reason for this behavior remains to be determined. LC-MS analysis of the doublet band suggests that both sub-bands present P2 (Supplementary Table 3). The P2 doublet is specific to recombinant wild-type protein and is not observed in the mutant $PanD_{127TRASC131}$ or in whole cell extracts (see Supplementary Fig. 6, Western blot analyses). The SDS-PAGE analyses show that the recombinant PanD protein preparations used for in vitro analyses in the current study presented largely the cleaved, active form of the enzyme.

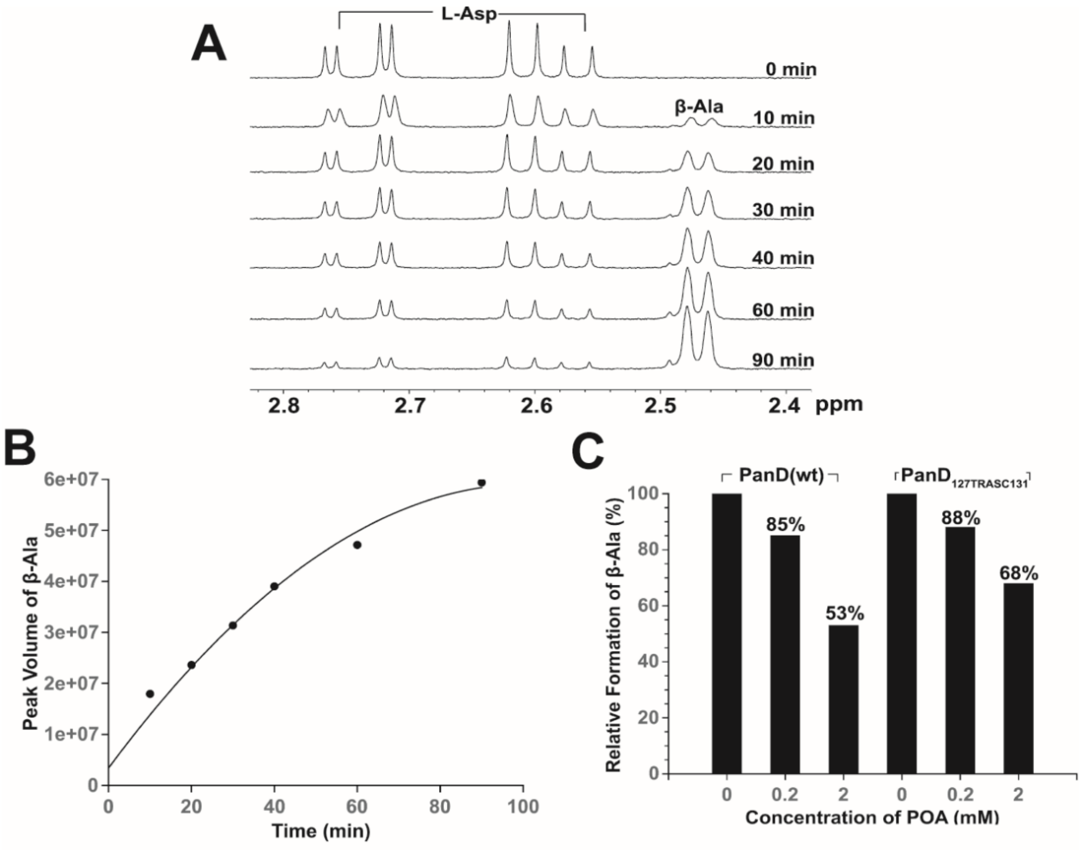


**Supplementary Fig. 2. Effect of POA on PanD enzymatic activity.** (A,B) Time-dependent conversion of L-Aspartate (2 mM) to β-Alanine (β-Ala) as determined by $^{1}H$ NMR after addition of 10 µM Mtb $PanD_{WT}$ in $D_2O$ at 298 K on a Bruker Avance 400 MHz spectrometer. Shown are (A) representative $^{1}H$ NMR spectra and (B) a plot of time-dependent β-Alanine formation at different incubation times by measuring the peak volume of converted L-Asp to β-Ala. At t = 40 min, approximately 50% conversion of L-aspartate to β-Alanine was observed as determined by integration of peak values. Thus, 40 min was chosen as a reference point within the linear range of PanD enzyme kinetics for subsequent enzyme assays. (C) Effect of POA on the formation of β-Ala by recombinant PanD wild-type ($PanD_{WT}$) and POA-resistant PanD mutant ($PanD_{127TRASC131}$) proteins, respectively, as determined by $^{1}H$ NMR. The plot shows the results of a representative experiment. NMR spectra for (C) are shown in Supplementary Fig. 3. Both recombinant proteins were purified according to [5]. SDS-PAGE analysis of the recombinant proteins is shown in Supplementary Fig. 1. Together, these data suggest that POA has only a minor inhibitory effect on the catalytic activity of PanD.

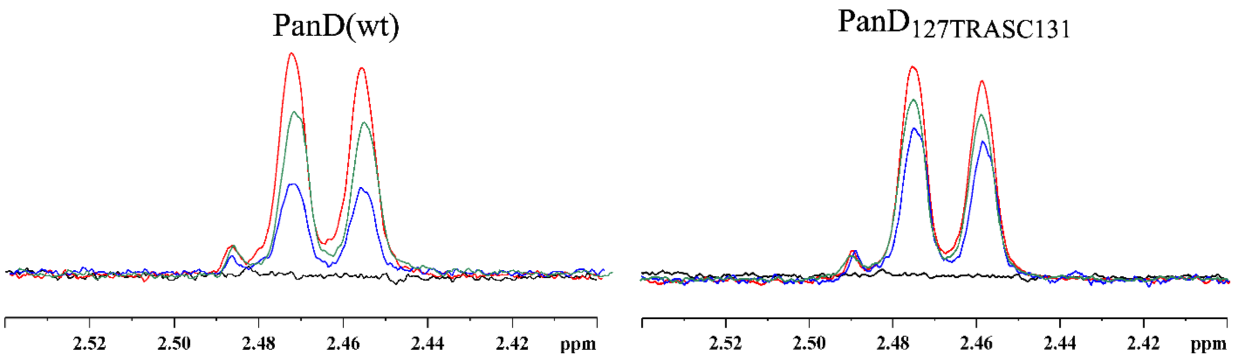


**Supplementary Fig. 3. Effect of POA on Mtb PanD enzyme activity as determined by $^{1}$H NMR.** Overlaid 1D NMR resonances of converted β-Alanine in $\text{PanD}_{\text{WT}}$ and POA-resistant C-terminal PanD mutant ($\text{PanD}_{\text{127TRASC131}}$) in the absence (*red*), and presence of 0.2 mM (*green*), or 2 mM (*blue*) POA after 40 min incubation at 298 K. The spectra in the absence of PanD enzymes are shown in black. The results of a representative experiment are shown.

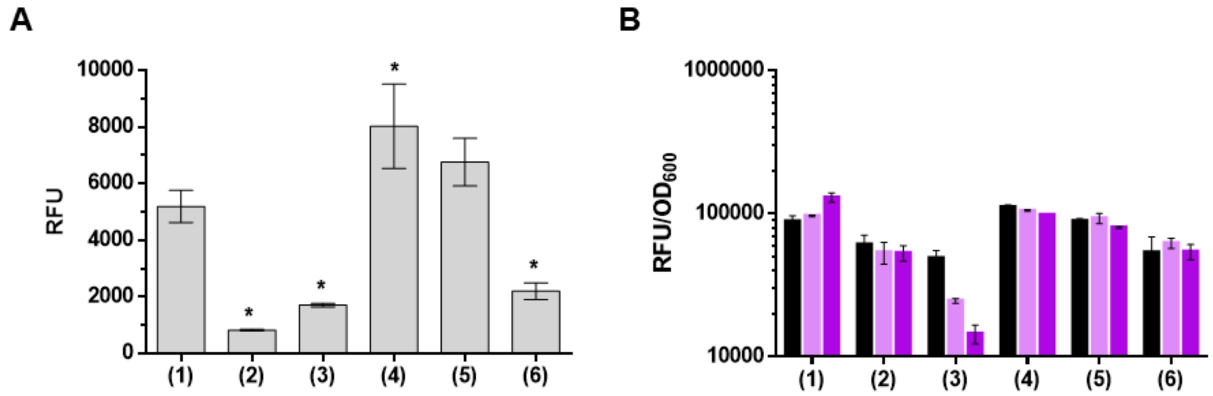


**Supplementary Fig. 4. Effect of POA treatment of *M. bovis* BCG on fluorescence levels of various PanD derivatives**. (A) Drug-free fluorescence levels of mid-log phase cultures were determined as a measure of intra-bacterial fusion protein level in *M. bovis* BCG carrying RFP-PanD reporter constructs described in Fig. 1A. All reporter strains were grown to mid-log phase and adjusted to a final $OD_{600} = 0.2$. (*) indicate RFU means that were significantly different from fluorescence levels of the native RFP control (construct (1)) in *M. bovis* BCG background at p-value $< 0.05$, one-way ANOVA multiple comparisons test, GraphPad Prism. (B) *M. bovis* BCG harboring various PanD derivatives fused translationally to constitutively expressed red fluorescence protein (RFP) as reporter (Fig. 1A) were treated with increasing doses of POA (light purple, 1 mM; dark purple, 2 mM). See Fig. 1A for structure of RFP-PanD fusion constructs #1-6. For all experiments, fluorescence (expressed as RFU) was used as a measure of intra-bacterial fusion protein level normalized to bacterial growth ($OD_{600}$), measured after incubation for 3 days at 310 K. For each strain and treatment, drug-free controls were included (black bars). Experiments were carried out two times independently in triplicates. Means and standard deviations from a representative experiment are shown. Taken together, these data suggest that the mechanism of action of POA in *M. bovis* BCG is the same as observed in Mtb.

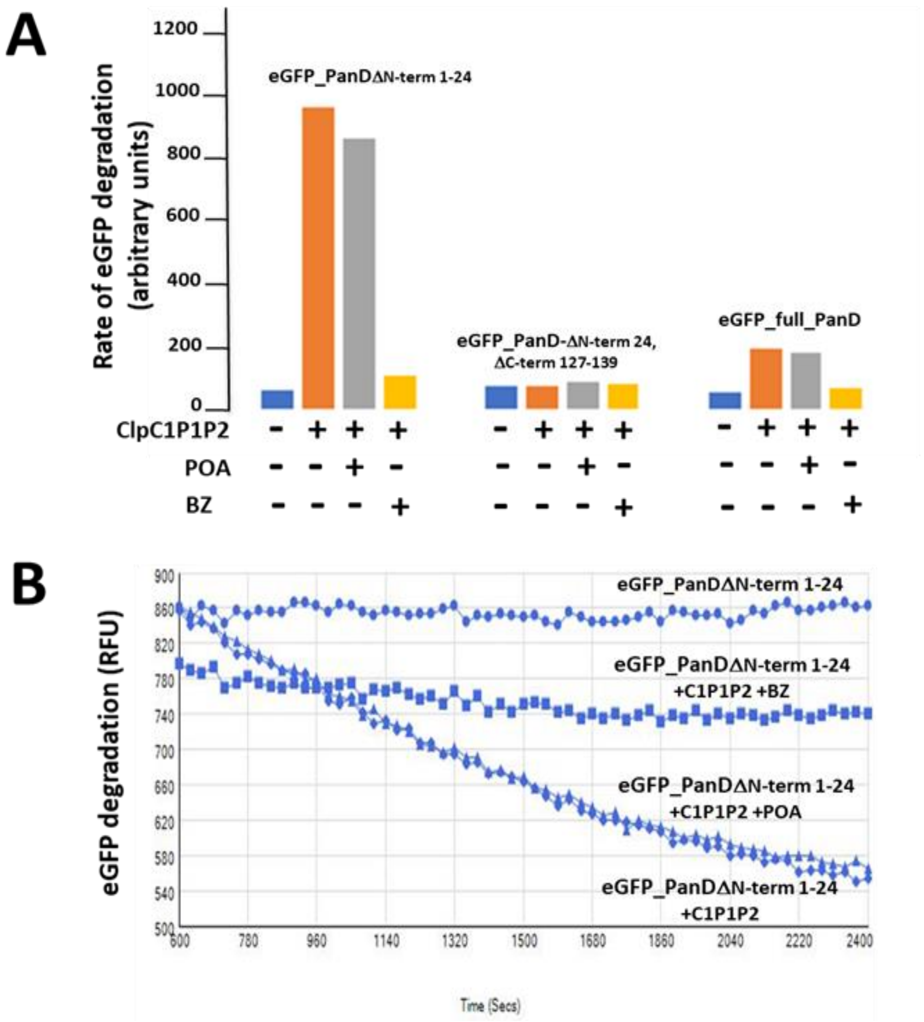


**Supplementary Fig. 5. *In vitro* degradation of PanD by ClpC1-ClpP.** (A) N-terminal truncated PanD containing its C-terminal 13 amino acid degradation tag (eGFP_PanD$_{\Delta\text{N-term1-24}}$), N-terminally truncated PanD without its C-terminal degradation tag (eGFP_PanD $_{\Delta\text{N-term1-24, }\Delta\text{C-term127-139}}$) and full-length PanD (eGFP_full_PanD) were subjected to an *in vitro* ClpC1-ClpP degradation assay as described [17]. PanD proteins were generated in a cell-free translation system and harbored eGFP fused at their N-termini to enable measurement of protein levels over time. Addition of ClpC1-ClpP (ClpC1-ClpP1P2) caused a strong decrease of fluorescence (i.e. degradation) for eGFP_PanD $_{\Delta\text{N-term1-24}}$. The fluorescence level of eGFP_PanD $_{\Delta\text{N-term1-24, }\Delta\text{C-term127-139}}$, i.e. eGFP_PanD $_{\Delta\text{N-term1-24}}$ lacking the C-terminal degradation tag, was not affected by ClpC1-ClpP, demonstrating that degradation depends on PanD's C-terminal tail. Degradation of eGFP_PanD $_{\Delta\text{N-term1-24}}$ was suppressed by the ClpP inhibitor bortezomib (BZ) demonstrating that the loss of fluorescence is due to proteolysis. As expected, POA had no effect on the rate of degradation as this version of PanD lacks the His21 residue required for drug binding [5]. Fluorescence of full-length eGFP_full_PanD was not affected by ClpC1-ClpP. The reason for the apparent lack of recognition and degradation of the full-length protein by the protease complex remains to be determined. Possible explanations include for instance differences in the quaternary structure of full-length PanD synthesized in a cell-free system when compared to protein produced *in vivo*. These differences may preclude access of the degradation tag by ClpC1-ClpP in the case of cell-free synthesized PanD. Additionally, auto-cleavage of full-length PanD may occur at Gly24-Ser25, potentially resulting in the detachment of reporter eGFP. In such a case, actual degradation would become non-detectable. BZ, bortezomib, concentration 100 µM; POA, pyrazinoic acid, concentration 1 mM. The experiments were carried out three times and a representative result is shown. (B)

Degradation of eGFP_PanD$_{\Delta N\text{-}term1\text{-}24}$ protein by ClpC1-ClpP in the absence or presence of bortezomib (BZ) or POA was followed continuously by measuring the change of eGFP fluorescence at 509 nm (ex 485 nm). RFU, relative fluorescence units.

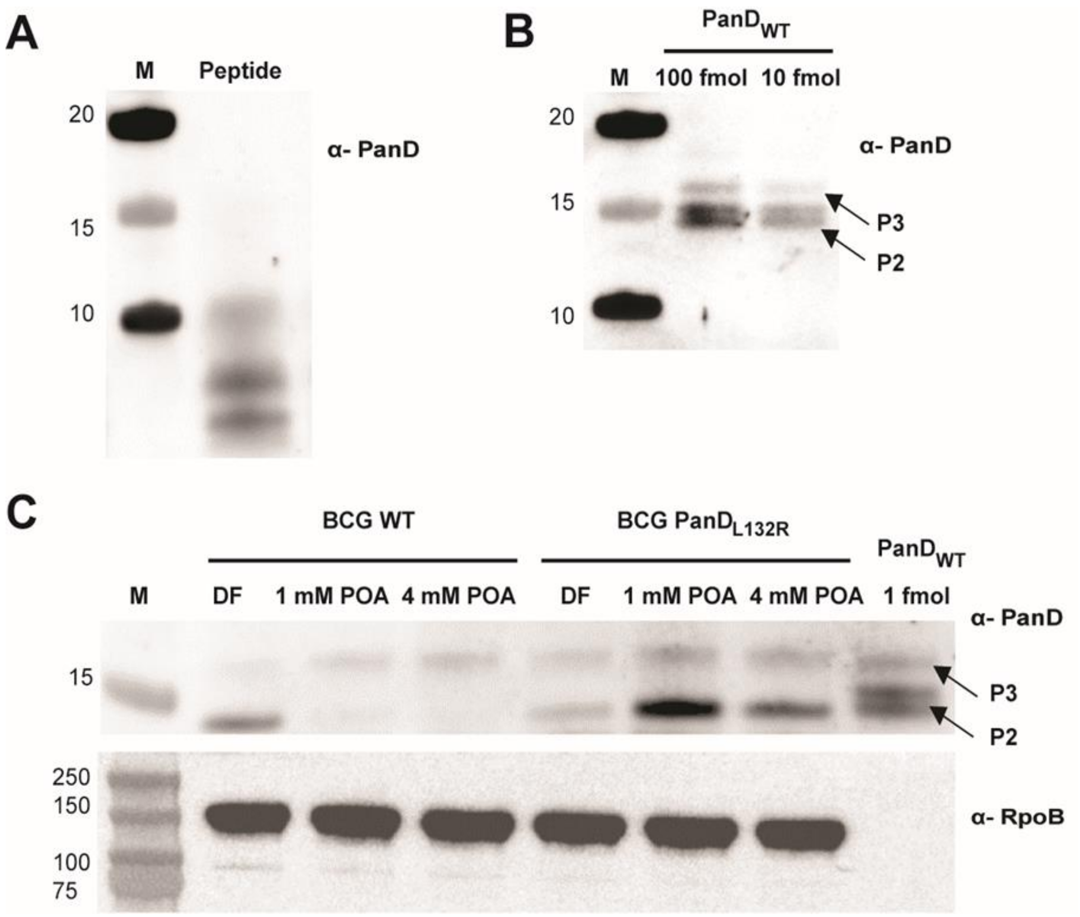


**Supplementary Fig. 6. Effect of POA treatment on PanD protein levels in wild-type and POA-resistant *panD*-mutant *M. bovis* BCG.** (A,B) Western blots of 1 µg PanD peptide IAYATMDDARARTY used as antigen, and 100 and 10 fmol purified recombinant PanD wild type protein ($PanD_{WT}$) respectively. P2, cleaved active form of PanD; P3 full-length proenzyme of PanD (Supplementary Fig. 1). Antiserum α-PanD against a PanD peptide was raised in rabbits and used to analyze protein levels of native PanD protein by Western blotting. Note: The cleaved N-terminal part of PanD (P1, Supplementary Fig. 1) is not visible on the blot as the PanD peptide used to raise the antiserum is located within the P2 portion of the protein. As observed in Coomassie blue stained SDS PAGE of purified recombinant PanD protein (Supplementary Fig. 1), the cleaved form of P2 migrates as a doublet with similar staining intensity. The reason for this behavior remains to be determined. LC-MS analysis of the doublet band suggests that both sub-bands represent P2 (Supplementary Fig. 1, Supplementary Table 3). The P2 doublet is specific to recombinant protein and is not observed in whole cell extracts (see (C)). (C) *M. bovis* BCG wild-type (BCG WT) and POA-resistant *M. bovis* BCG $PanD_{L132R}$ harboring a Leu132Arg mutation in its C-terminal tail [7] were either not treated with POA (drug free, DF) or treated with 1 or 4 mM POA for 24 h. 10 µg of respective total protein extracts were subjected to Western blot analyses. Upper panel: probing with α-PanD. $PanD_{WT}$, 1 fmol of recombinant PanD wild type protein was included as molecular weight marker. Lower panel: probing of the blot showed in the upper panel with antiserum against mycobacterial RNA polymerase subunit RpoB (α-RpoB) to show equal loading. M, Marker with molecular weights (kilodalton) are indicated. Experiments were repeated 3 times independently and representative results are shown. Taken together, these data show that treatment with POA causes a reduction of the intra-bacterial level of wild-type PanD but does not affect the intra-bacterial level of a non-POA binding resistant mutant of PanD.

Note: Consistent with the overall low intra-bacterial levels of PanD indicated by our Western blotting experiments, PanD was not detectable in previous proteome analyses (<1 fmol/µg), whereas all other CoA pathway enzymes could be quantified (1-6 fmol/µg) [29]. We confirmed this finding by shotgun proteomic analyses of mycobacterial whole cell extracts in which all CoA pathway enzymes except PanD were readily detectable, including the pantothenate synthetase PanC (Supplementary Table 4).

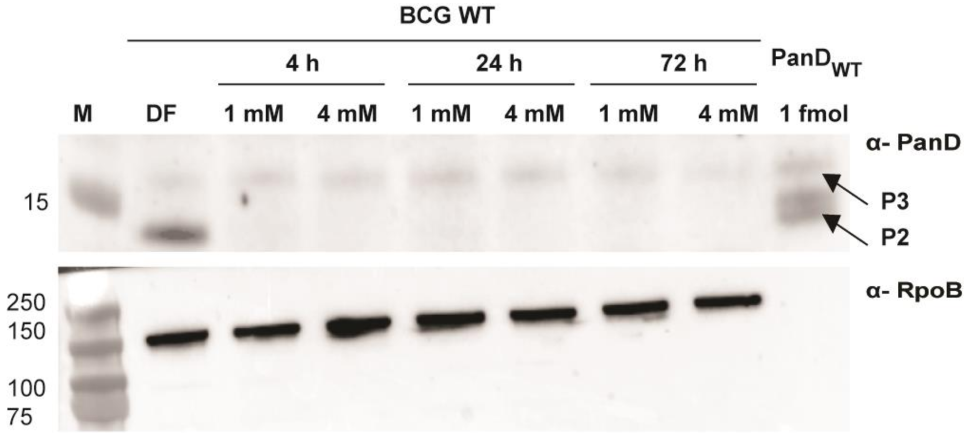


**Supplementary Fig. 7. Time kinetics of POA-induced PanD degradation in *M. bovis* BCG.** *M. bovis* BCG wild-type (BCG WT) was treated with 1 mM or 4 mM POA and cells were harvested at 4 h, 24 h or 72 h after treatment for extraction of total protein. Drug-free (DF) controls were harvested at the start of the experiment (t = 0). 10 µg of respective total protein extracts were subjected to Western blot analyses. Upper panel: probing with α-PanD. PanD$_{WT}$, 1 fmol of recombinant PanD wild type protein was included as molecular weight marker. Lower panel: probing of blot showed in the upper panel with antiserum against mycobacterial RNA polymerase subunit RpoB (α-RpoB) to show equal loading. M, Marker with molecular weights (kilodalton) are indicated. Experiments were repeated 3 times independently and representative results are shown. These results show that POA treatment causes a rapid (4 h) and sustained (72 h) reduction of intra-bacterial PanD levels.

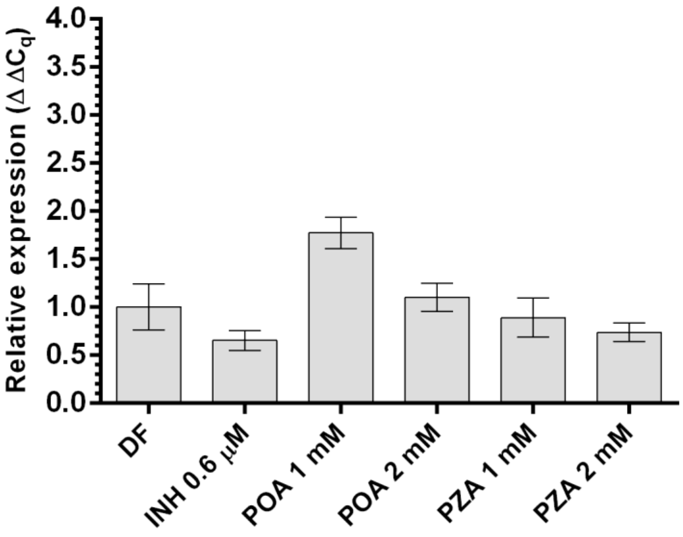


**Supplementary Fig. 8. *panD* mRNA levels in wild-type Mtb upon drug treatment.** Transcript levels were measured from mid-log phase cultures (equivalent of 20 mL of $OD_{600}$ = 0.4 were collected for each sample), adjusted to $OD_{600}$ = 0.2 and incubated with or without drug for 24 h at 310 K. Drug-free (DF) controls were obtained at t = 0 h. INH 0.6 µM corresponds to 2 x Mtb $MIC_{50}$. Primer sequences can be found in Supplementary Table 2. Relative expression (quantification cycle [ΔΔCq]) was calculated as described previously [30] by using 16S RNA as the reference. The experiment was repeated two times independently. Mean and SD from technical triplicates of a representative experiment are shown. Means were found not to be significantly different from DF controls at p-value < 0.05 (*), one-way ANOVA multiple comparisons and Dunn's posttest, GraphPad Prism. These data show that treatment of bacteria with POA or PZA does not affect the mRNA level of *panD*.

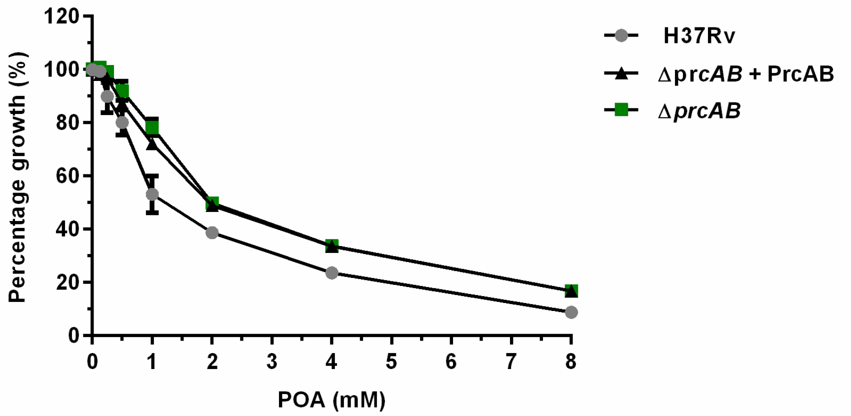


**Supplementary Fig. 9. Effect of deletion of the mycobacterial proteasome genes *prcAB* on the growth inhibitory activity of POA.** The effect of increasing POA concentrations on growth inhibition of *M. tuberculosis* H37Rv, *M. tuberculosis* Δ*prcAB* and the *prcAB* complemented mutant strains [18], respectively. The experiment was carried out two times independently. Mean and standard deviations from a representative experiment are shown. These data suggest that the mycobacterial proteasome is not involved in the mechanism of action of POA.

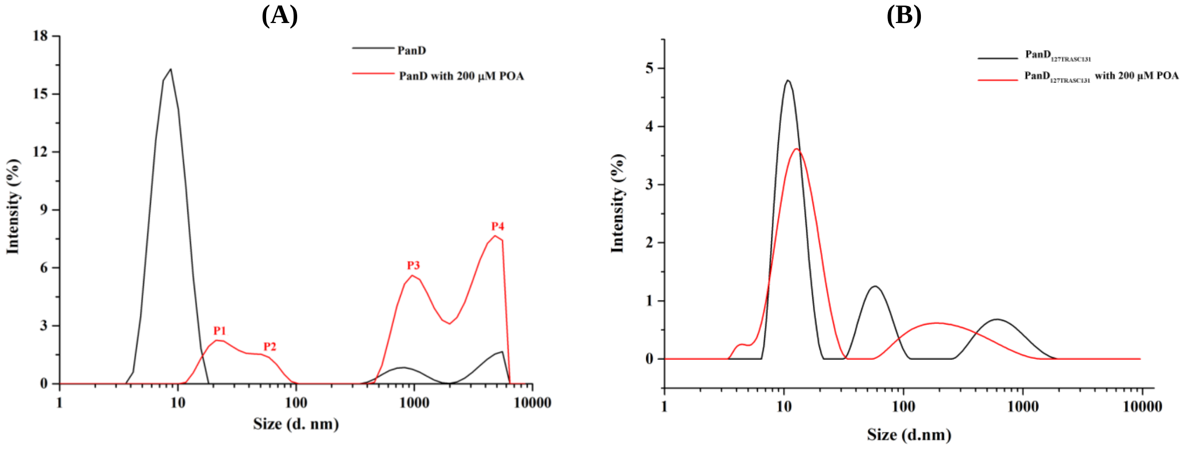


**Supplementary Fig. 10. Effect of POA on PanD$_{WT}$ and resistant mutant, non-POA binding PanD$_{127TRASC131}$ as determined by dynamic light scattering.** (A) The distribution of the intensity of dynamic light scattering by PanD$_{WT}$ without drug (*black*) and PanD$_{WT}$ with 200 µM POA (*red*) as a function of particle diameter (*d* in nm) shows that PanD$_{WT}$ transforms into higher oligomers (P1-4) upon addition of POA with calculated hydrodynamic diameters of 21.04 ± 7.22 nm, 55.04 ± 6.35 nm, 955.4 ± 106.9 nm and 4801 ± 1134 nm, respectively. The estimated intensity of peak 1 to 4 were 11%, 8%, 38%, and 43%, respectively. (B) In contrast, the PanD$_{127TRASC131}$ mutant retained its prominent peak at around 11.5 nm in the presence (*red*) of 200 µM POA (drug free control: *black*). All experiments were repeated three times yielding the same results. These data show that POA causes oligomer formation of wild-type PanD but not of a POA resistant mutant version of the protein. These results suggest that the drug does not cause protein aggregation *per se*.

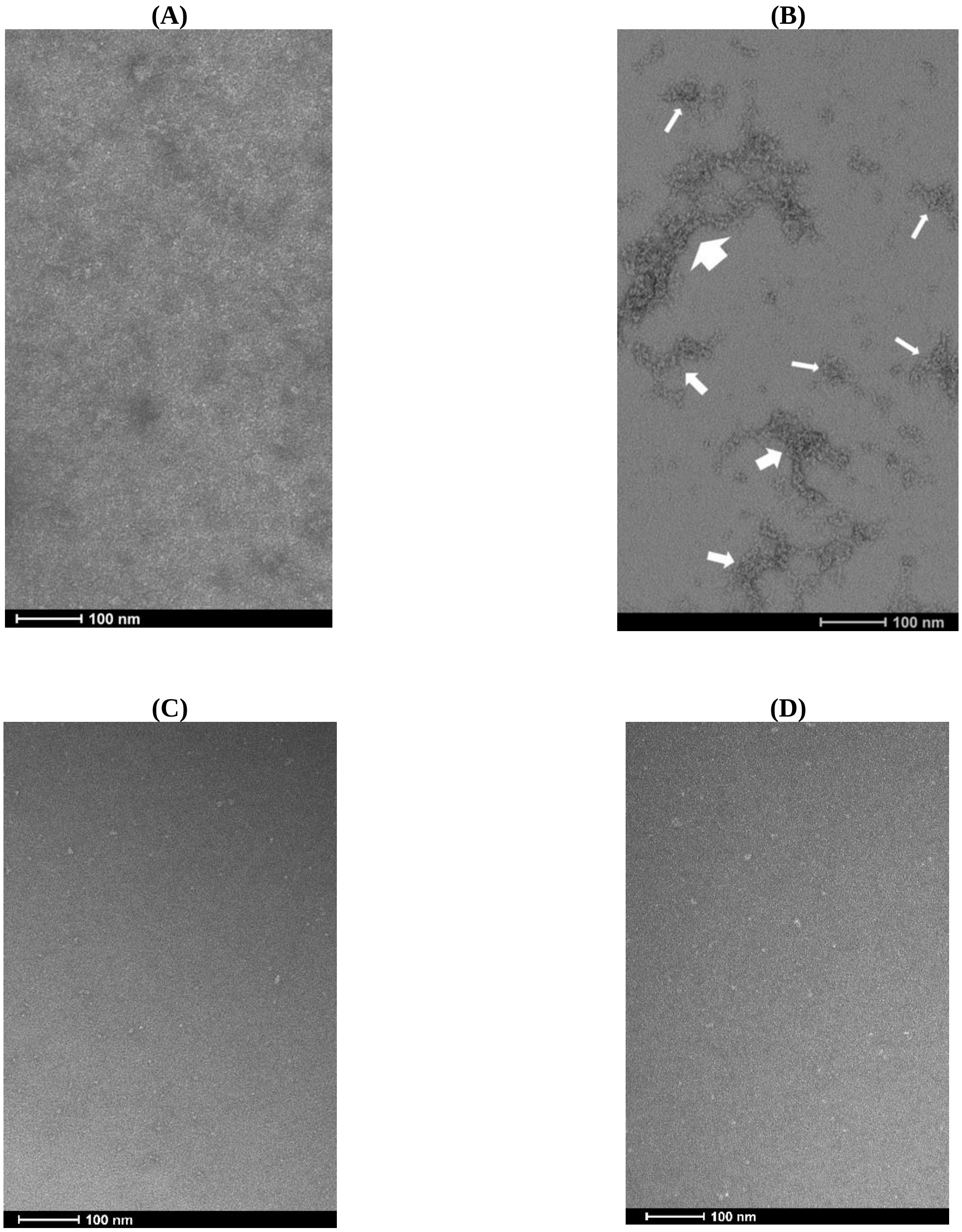


**Supplementary Fig. 11. Effect of POA on PanD$_{WT}$ and resistant mutant, non-POA binding PanD$_{127TRASC131}$ as visualized by electron microscopy.** Electron-micrograph of negatively-stained PanD$_{WT}$ in the absence (A) and presence of 200 µM POA (B). The comparison visualizes oligomer formation upon drug-binding to PanD$_{WT}$. The different sized white arrows indicate the different sizes of oligomers formed. (C-D) Control experiment showing PanD$_{127TRASC131}$ mutant protein in absence (C) and presence (D) of 200 µM POA, demonstrating that the formation of aggregates by POA is specific to the wild-type protein. Scale bars: 100 nm. These data show that POA causes oligomer formation of wild-type PanD but not of a POA-resistant mutant version of the protein. These results are consistent with the corresponding results from dynamic light scattering experiments shown in Supplementary Fig. 10.

**(A)**

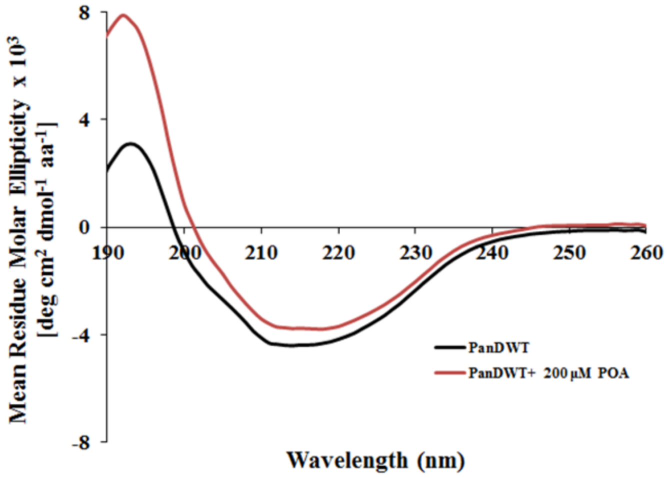


**(B)**

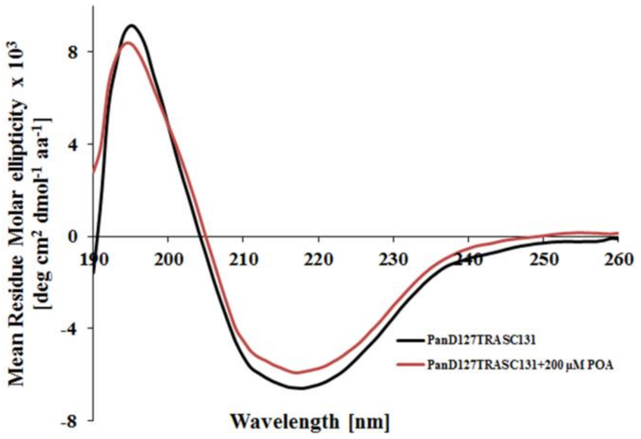


**Supplementary Fig. 12. Effect of POA on PanD$_{WT}$ and resistant mutant, non-POA binding PanD$_{127TRASC131}$ as determined by circular dichroism spectroscopy.** (A) Far-UV CD spectra of PanD$_{WT}$ without drug (*black*) and PanD$_{WT}$ with 200 µM POA (*red*). Two independent measurements were carried out and three spectra per sample were collected, yielding the same results. The far UV spectrum of PanD$_{WT}$ revealed a peak at 195 nm and a broad minimum at 218 nm indicative of a significant presence of β-sheets. The CD spectrum of drug-free PanD suggests a secondary structural content of 4% α-helices and 37.8% β-sheets, consistent with the secondary structural content determined by the crystallographic structure of PanD (PDB ID: 2C45). The presence of 200 µM POA caused an increase of mean residual molar ellipticity and decrease in breadth of the spectrum. (B) The far-UV CD spectra of the PanD$_{127TRASC131}$ mutant protein in the absence (*black*) and presence (*red*) of 200 µM POA. The CD spectra of the PanD$_{127TRASC131}$ mutant in the presence and absence of 200 µM POA did not show the drastic effect at the 195 nm peak as observed for PanD$_{WT}$ shown in A. No change in the broader range from 190-210 nm was observed for the mutant protein in the presence of the drug. Only a minor decrease of mean residual molar ellipticity was detected at the 218 nm minima in the mutant spectrum after addition of 200 µM POA. These data show that POA causes a drastic change of the CD spectrum of wild-type PanD but has only minor effects on the CD spectrum of a POA-resistant mutant version of the protein.

**Supplementary Table 1.** Fluorescent protein fusion plasmids and primers used in this study

| Plasmid Name | Backbone plasmid (digested with) | Template DNA for PCR in this study | Inserted PCR-amplified DNA fragments | | Reference (if any) |
|---|---|---|---|---|---|
| | | | Primer Name | Primer Sequence (5'->3') | |
| (1) RFP | pMV262 (BamHI-EcoRI) | Not applicable | mCh-F(BamHI)<br>mCh-R(EcoRI) | ccgggatccATGGTGAGCAAGGGCGAGG<br>ccggaattcCTACTTGTACAGCTCGTCCAT | *(14)* |
| (2) RFP-C-terminal | pMV262 (EcoRI-HindIII) | *Mtb* H37Rv | PanD FORWARD EcoRI_T<br>PanD REVERSE HindIII_T | ggaattcAACGCGGGCGAG<br>cccaagcttCTATCCCACACCG | This study |
| (3) RFP-PanD | pMV262 (EcoRI-HindIII) | *Mtb* H37Rv | PanD FORWARD EcoRI<br>PanD REVERSE HindIII | ccggaattcATGTTACGGACG<br>cccaagcttCTATCCCACACC | This study |
| (4) RFP-PanD no C-terminal | pMV262 (EcoRI-HindIII) | *Mtb* H37Rv | PanD FORWARD EcoRI<br>PanD w/o C-term REVERSE HindIII | ccggaattcATGTTACGGACG<br>cccaagcttCTATTCGGGCAC | This study |
| (6) RFP-PanD$_{\mathbf{L136R}}$ | pMV262 (EcoRI-HindIII) | *M. bovis* BCG POA1.1* | PanD FORWARD EcoRI | ccggaattcATGTTACGGACG<br>cccaagcttCTATCCCACACC | This study |

| | | | | | |
|---|---|---|---|---|---|
| | | | PanD REVERSE HindIII | | |
| (7) RFP-PanD$_{\mathbf{H21R}}$ | pMV262 (EcoRI-HindIII) | *M. bovis* BCG POA1.3* | PanD FORWARD EcoRI<br><br>PanD REVERSE HindIII | ccggaattcATGTTACGGACG<br><br>cccaagcttCTATCCCACACC | This study |
| pET26b_eGFP! | pET26b+ (NdeI-HindIII) | Not applicable | JH-306<br><br>JH-311 | aactttaagaaggagatatacatatgtcgaagggcgaggagctg<br><br>tgctcgagtgcggccgcaagcttatttgtacagctcgtccatgccca | This study |
| pET26b_eGFP_<br><br>full_PanD! | pET26b+ (NdeI-HindIII) | *Mtb* H37Rv | JH-PAND_1<br><br>JH-PAND_2 | tacaaaggatctagcggatccagtttacggacgatgctgaagtcg<br><br>tgctcgagtgcggccgcaagcttatcccacaccgagccggggg | This study |
| | | pET26b_eGFP | JH-306<br><br>JH-307 | aactttaagaaggagatatacatatgtcgaagggcgaggagctg<br><br>actggatccgctagatcctttgtacagctcgtccatgcc | |
| pET21a_eGFP_PanD$_{\Delta\text{N-term1-24}}$# | pET21a (NdeI-HindIII) | *Mtb* H37Rv | JH_PAND_11<br><br>JH-PAND_2 | tacaaaggatctagcggatccagttcggtgaccatcgatgccg<br><br>tgctcgagtgcggccgcaagcttatcccacaccgagccggggg | This study |
| | | pET26b_eGFP | JH-306<br><br>JH-307 | aactttaagaaggagatatacatatgtcgaagggcgaggagctg<br><br>actggatccgctagatcctttgtacagctcgtccatgcc | |
| pET21a_eGFP_PanD$_{\Delta\text{N-term1-24, C-term 127-139}}$# | pET21a (NdeI-HindIII) | *Mtb* H37Rv | JH_PAND_11<br><br>JH-PAND_7 | tacaaaggatctagcggatccagttcggtgaccatcgatgccg<br><br>tgctcgagtgcggccgcaagcttattcgggcacaaatgccggat | This study |
| | | pET26b_eGFP | JH-306<br><br>JH-307 | aactttaagaaggagatatacatatgtcgaagggcgaggagctg<br><br>actggatccgctagatcctttgtacagctcgtccatgcc | |

* *M. bovis* BCG POA1.1 and *M. bovis* BCG POA1.3 were isolated and described in [7].

Numbered plasmids 1-7: used for *in vivo* studies. Non-numbered plasmids: used for *in vitro* translation.

[!] *in vitro* translation vector maintained in *E. coli* DH5α with Kanamycin selection.

[#] *in vitro* translation vector maintained in *E. coli* TOP10 with Ampicillin selection.

**Supplementary Table 2**. Primers used for qRT-PCR

| **Target** | **Primer** | **Primer sequence** | **Amplicon (bp)** | **Reference** |
|---|---|---|---|---|
| *16S* | Fwd<br>Rev | ATGACGGCCTTCGGGTTGTAA<br>CGGCTGCTGGCACGTAGTTG | 160 | *(14)* |
| *panD* | Fwd<br>Rev | TACGGACGATGCTGAAGTCG<br>CGATGGTTACCTGTTCGCCT | 135 | This study |
